# Physiological activation of the nephron central command drives endogenous kidney tissue regeneration

**DOI:** 10.1101/2021.12.07.471692

**Authors:** Georgina Gyarmati, Urvi Nikhil Shroff, Anne Riquier-Brison, Dorinne Desposito, Wenjun Ju, Audrey Izuhara, Sachin Deepak, James L Burford, Hiroyuki Kadoya, Yibu Chen, Markus M. Rinschen, Nariman Ahmadi, Lester Lau, Inderbir S. Gill, Matthias Kretzler, János Peti-Peterdi

## Abstract

Tissue regeneration is limited in several organs including the kidney, contributing to the high prevalence of kidney disease globally. However, evolutionary and physiological adaptive responses and the presence of renal progenitor cells suggest existing remodeling capacity. This study uncovered a novel endogenous tissue remodeling mechanism in the kidney that is activated by the loss of body fluid and salt and involves a unique niche of chief cells called macula densa (MD) that control resident progenitor cells via secreted angiogenic, growth and extracellular matrix remodeling factors, cytokines and chemokines. Serial intravital imaging, MD Wnt mouse models and transcriptome analysis provide functional and molecular characterization of this newly identified MD program for kidney regeneration complemented with human and therapeutic translation. The concept that chief cells responding to organ-specific physiological inputs control local progenitors and direct them to remodel or repair tissues may be applicable to other organs and diverse tissue regenerative therapeutic strategies.

## Introduction

Regeneration of most tissues and organs in mammalians, particularly in humans is limited (*1*), and the kidney is a prime example (*2*). Despite recent advances in regrowing or repairing kidney tissues, chronic kidney disease remains a major global health issue and its incidence continues to rise (*2*). Improved understanding of endogenous kidney tissue repair and identification of the key cellular and molecular targets are critically important for the development of specific, mechanism-based regenerative therapies.

Physiological adaptation to maintain functional homeostasis is a known driver of physiological regeneration in multiple organisms and mammalian tissues (*1*). Physiological signals associated with either loss or gain of organ function, such as fasting for the nervous, endocrine and digestive systems (*3, 4*) and mechanical force for the skeletal system (*5*) can trigger potent regenerative responses. Additional clues regarding endogenous regenerative mechanisms come from adaptive evolution. Conservation of body fluid and salt was a major driver of evolution that brought about the highly efficient and complex mammalian nephron compared to the primitive structure in fish (*6*). The loss of neo-nephrogenesis capacity in birds and mammals coincided with the appearance of a cell-based regeneration strategy and the differentiation of unique, specialized nephron cell types (*6*) including the macula densa (MD). MD cells are chief salt sensors in the kidney and function as a neuron-like central command for the nephron (*7*).

Here we addressed the hypothesis that MD cells control progenitor cell-mediated endogenous kidney regeneration that can be activated by the loss of body fluid and salt, which is the primary, organ-specific and evolution-driven physiological signal for the kidney.

## Results

### Live tracking and MD-centric pattern of endogenous tissue remodeling

To establish the dynamics and pattern of endogenous kidney tissue remodeling with single-cell resolution in the intact living kidney, unbiased tracking of the same tissue volume of kidney cortex over several days and weeks was performed using a combination of serial intravital multiphoton microscopy (MPM)(*8, 9*) and genetic cell fate tracking. In contrast to the lack of effect of pathological injury (ischemia-reperfusion) (Fig. S1A), the loss of body fluid and salt as a MD-activating physiological stimulus using treatment with low-salt (LS) diet+angiotensin converting enzyme inhibitor enalapril (ACEi) for two weeks caused substantial recruitment of mesenchymal and endothelial precursor cells in Ng2-tdTomato and Cdh5-Confetti mice (*10*), respectively (Fig. 1A and Supplement Movie 1). Importantly, the geometrical epicenter (highest cell density) of both mesenchymal and endothelial cell recruitment was the base of MD cells at the glomerular vascular pole in each nephron (Fig. 1A). Other physiology-based stimuli that are known to trigger MD cell sensing of low salt (the diuretic furosemide, low salt diet or ACEi alone) caused similar, but less robust responses (Fig. S1A). Newly recruited Ng2^+^ cells differentiated to multiple cell fates including vascular smooth muscle, renin, mesangial, parietal epithelial and proximal tubule cells and podocytes (Fig. S1B). Alternative genetic strategies to track mesenchymal cell lineages (Ren1d-Confetti and Foxd1-tdTomato mice) produced similar results (Fig. S1C). Both Cdh5^+^ and Ren^+^ cells produced clonal remodeling of the vasculature, interstitium and glomerulus closest to the MD (Figs. 1A and S1C), further suggesting the presence of mesenchymal and endothelial progenitor cells at the glomerular vascular pole. LS+ACEi-induced Ng2^+^ and Cdh5^+^ cell recruitment to the glomerular vascular pole was blocked by pharmacological inhibition of the known MD-specific markers cyclooxygenase-2 (Cox-2) or neuronal nitric oxide synthase (Nos1), suggesting the key role of MD cells in this tissue remodeling process (Fig. 1B). Fate tracking of MD cells for extended periods of time (<6 months) in either control or LS+ACEi stimulation conditions using inducible MD-GFP mice found no GFP-labeled cells outside of the MD area (data not shown).

**Figure 1.**
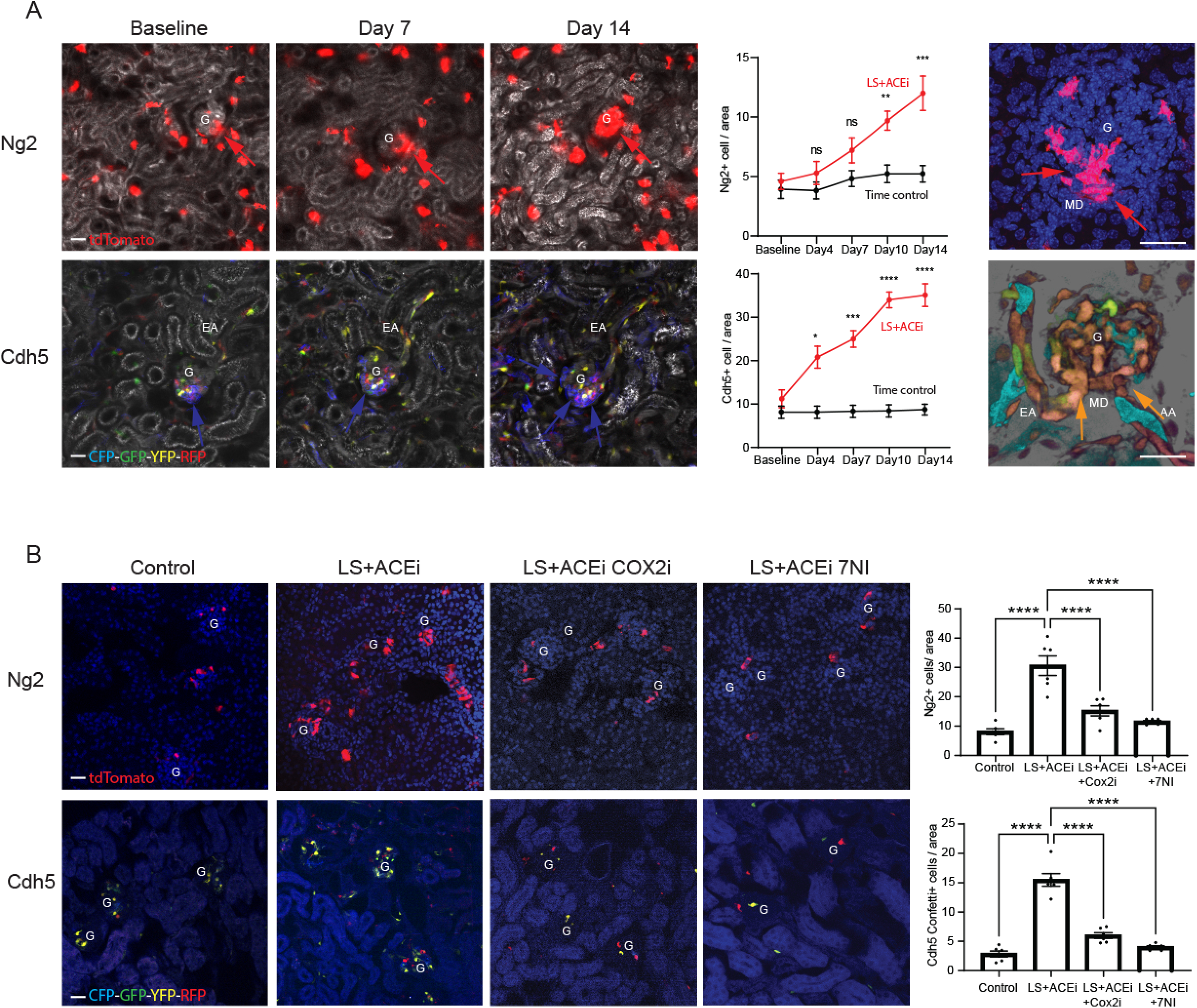
Evaluation of mesenchymal (Ng2-tdTomato, red) and endothelial (Cdh5-Confetti, multicolor) precursor cell-mediated endogenous kidney tissue remodeling by serial intravital MPM imaging of the same glomeruli/kidney over several days-weeks (A) or by conventional histological analysis (B). **(A)** Representative *in vivo* MPM maximum projection images (left) and the number of Ng2^+^ or Cdh5^+^ cells per glomerular area (center) of the same kidney cortex area/volume visualized through an abdominal imaging window at the indicated time points or in a magnified single glomerulus (right, on Day14). Responses to low-salt (LS) diet+ACEi (all panels) or timed control (center) are shown. Note the low cell number and random distribution at baseline, but high cell number specifically at the glomerular vascular pole (red and blue arrows) and the Cdh5^+^ clones (CFP/blue (left) and YFP/RFP orange (right)) among all 10 different CFP/GFP/YFP/RFP Confetti color combinations. Plasma was labeled with Alexa594-Albumin (greyscale). The z-stacks of same preparations are shown in Supplement Movie 1. G: glomerulus, AA/EA: afferent/efferent arteriole, MD: macula densa. **(B)** The effects of LS diet+ACEi for 10 days with or without selective Cox-2 inhibition with SC58236 or Nos1 inhibition with 7-NI treatment, or 10 days timed control on Ng2^+^ and Cdh5^+^ cell number per glomerular area analyzed on frozen tissue sections. Nuclei are labeled blue with DAPI. Scale bar is 50 μm for all panels. Ns: not significant, *: p<0.05-****p<0.0001, data are mean ± SEM, n=5 glomeruli averaged for n=6 mice each group.

### The molecular profile of the MD cell program for tissue regeneration

We next aimed to identify the molecular signature of the MD cell tissue remodeling program in both control and physiological activation states using MD and control cell isolation from MD-GFP mouse kidneys as established and validated recently (*11*), followed by bulk and single-cell RNA sequencing and transcriptomic analysis (Fig. 2A). A total of 28,000 MD cells (representing a very minor fraction, ∼0.2% of the total kidney cortical cell population) and 50,000 control cells from adjacent tubule segments were isolated from the cortex of freshly digested MD-GFP kidneys (n=2 mice for MD and n=4 mice for control cells from each condition) for bulk RNA isolation and transcriptome analysis. For single-cell RNA sequencing and transcriptome analysis, 894 and 1296 MD cells were analyzed from control and LS-induced conditions, respectively, each from a single MD-GFP mouse. Single-cell transcriptome analysis identified angiogenesis, cell movement, quantity and migration of cells and vascular endothelial cells as the major biological activities of MD cells (Fig. 2B). Unsupervised graph-based clustering and UMAP visualization using Partek Flow revealed five subtypes of MD cells under physiological activation (LS) condition (MD1-5, Fig. 2C) confirming our recent finding under control condition (*7*). Certain MD cell clusters showed high expression of specific growth and transcription factors and chemokines (*Fabp3, Egf, Foxq1, Cxcl12* for MD3) and angiogenic factors (*Vash2, Pamr1, Vegfa, Nov* for MD5), suggesting MD cell heterogeneity regarding their tissue remodeling control functions. In contrast, all MD clusters were similarly enriched in other secreted cytokines, growth and extracellular matrix (ECM) and Wnt signaling factors (*Bmp3, Fgf9, Spp1, Wnt10a, Sfrp2, Tcf4*) that have well-known roles in controlling progenitor cells and tissue remodeling. Bulk RNA-based MD transcriptomic analysis further confirmed the MD-specific expression of several angiogenic, cell migration and patterning, growth, ECM remodeling and transcription factors. The expression of these factors was upregulated in physiological activation (LS) condition compared to control (Fig. 2D and Supplement Table 1). The highly MD-specific expression of a selection of these genes was validated and translated to the human kidney on the protein level (Fig. 2E).

**Figure 2.**
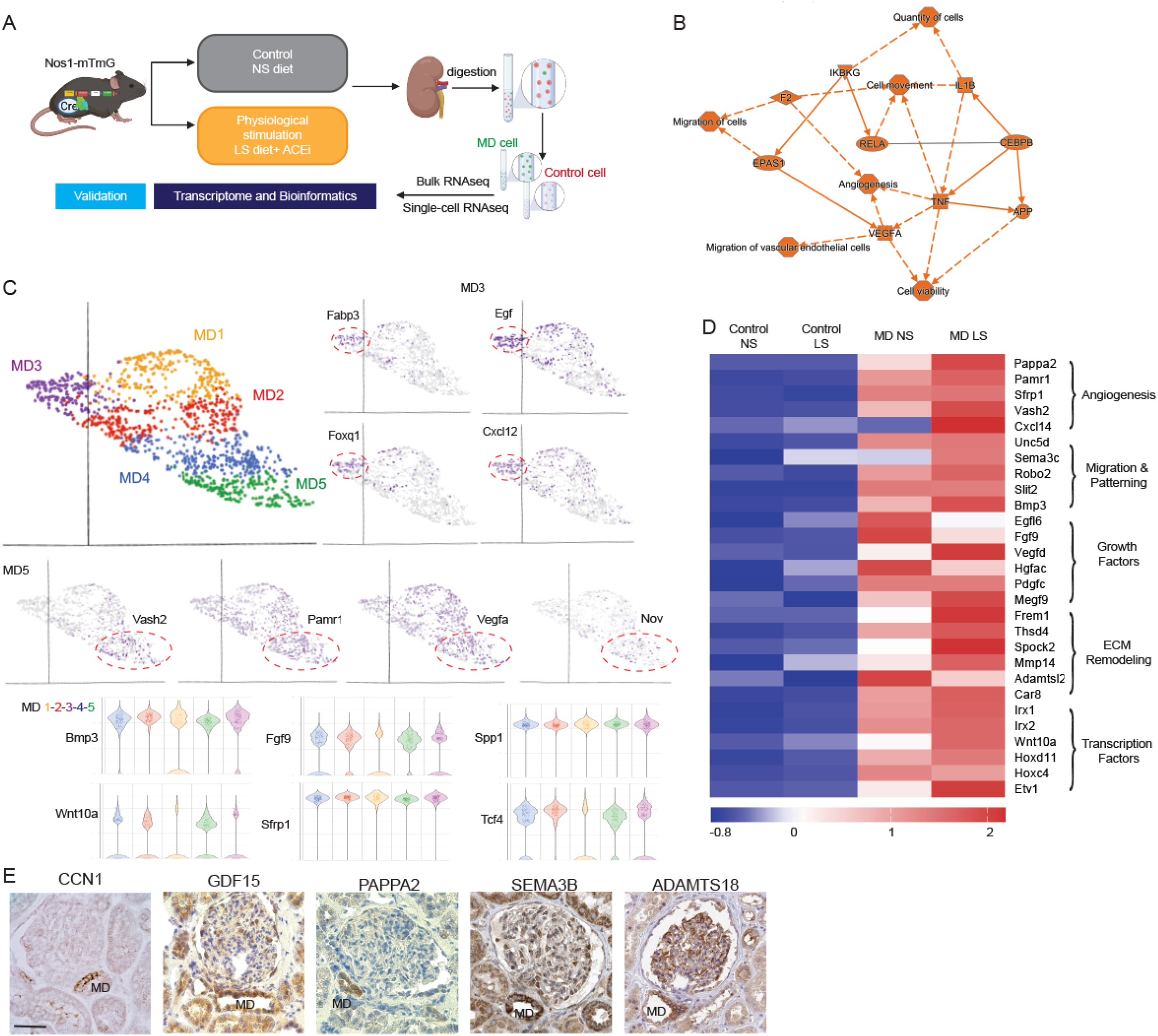
Tissue regenerative profile of the MD cell transcriptome. **(A)** Workflow of the MD cell transcriptome analysis. MD (green) and control (red) cells were freshly harvested and digested into single cell suspensions from MD-GFP (Nos1-mTmG) mouse kidneys in control and physiological activation states, and cells were analyzed and sorted based on their fluorescence protein expression (GFP for MD, tdTomato for control cells) followed by either bulk or single-cell RNA isolation and sequencing. **(B)** Graphical summary of MD single-cell transcriptome analysis. The top activated (indicated by orange, positive z-score) biological activities, pathways and genes are listed based on unbiased IPA analysis. **(C)** UMAP visualization (top) of integrated single-cell transcriptomic analysis of MD cells (n = 1296 MD cells from single MD-GFP mouse in LS condition). Five cell clusters (MD1-5) were identified based on graph-based analysis in Partek Flow. Cluster-specific expression of top enriched genes in MD3 (Fabp3, Egf, Foxq1, Cxcl12) and MD5 clusters (Vash2, Pamr1, Vegfa, Nov). Violin plot (bottom) of the expression of highly enriched MD genes in all five clusters (MD1-5). **(D)** Heat map of the expression of top enriched MD-specific genes in MD vs. control cells and in control and physiological stimulation (LS) conditions based on bulk RNA analysis. Genes were grouped into five categories as indicated according to their biological function. Scale indicates z-score values. **(E)** Immunohistochemistry validation of the expression of top enriched MD-specific genes in the human kidney. Data are from the Human Protein Atlas if indicated. Scale bar is 50 μm for all panels.

### MD cell Wnt signaling controls glomerular remodeling

Wnt/β-catenin signaling is one of the key regulators of tissue maintenance and regeneration (*12*), and many of the newly identified top MD-enriched genes (Fig. 2) are known Wnt targets. In addition, new results show that among all renal cell types in the renal cortex, MD cells have the highest Wnt activity (Fig. S2A). Therefore, we next addressed the kidney tissue remodeling role of Wnt signaling specifically in MD cells using inducible genetic MD Wnt loss-of-function (MD-Wnt^lof^) and gain-of-function (MD-Wnt^gof^) mouse models (Fig. 3A) that were generated based on Cre/lox-mediated loss or accumulation of β-catenin in MD cells, respectively. Altered MD Wnt activity in these mouse models were validated by the increased or decreased MD-specific genetic WntGFP reporter expression, and mRNA expression of the Wnt target Axin2 in MD-Wnt^gof^ or MD-Wnt^lof^ mouse kidneys, respectively (Fig. S2B). Blood pressure, kidney and body weight, and albuminuria were normal in these mouse models (Fig. S2B). However, the induction of MD-Wnt^gof^ in adult mice resulted in enlarged, hypercellular cortical glomeruli and increased glomerular filtration rate (GFR) within 6-8 weeks (Fig. 3B). In contrast, the induction of MD-Wnt^lof^ resulted in smaller, hypocellular glomeruli and reduced GFR (Fig. 3B). The number of WT1^+^ mesenchymal progenitor cells/podocytes and CD34^+^ endothelial precursor cells (alternatively Meis2^+^/Ren^+^ endothelial/mesenchymal cells, respectively, Fig. S2C) at the glomerular vascular pole increased in a MD-centric pattern in MD-Wnt^gof^ mice, while these cell numbers were reduced or unchanged in MD-Wnt^lof^ mice (Fig. 3C). Regarding the MD-specific synthesis of secreted, paracrine acting tissue remodeling factors, MD-Wnt^gof^ signaling promoted global MD protein synthesis and specifically their angiogenic output, including Ccn1 and other angiogenesis promoting factors, including Sema3C that are known Wnt target genes (*13*)(Fig. 3D-E). Altogether, these data strongly suggested that MD cells via Wnt signaling are major regulators of kidney tissue maintenance and have the capacity to quickly remodel the renal vasculature, interstitium and glomerulus both structurally and functionally when activated.

**Figure 3.**
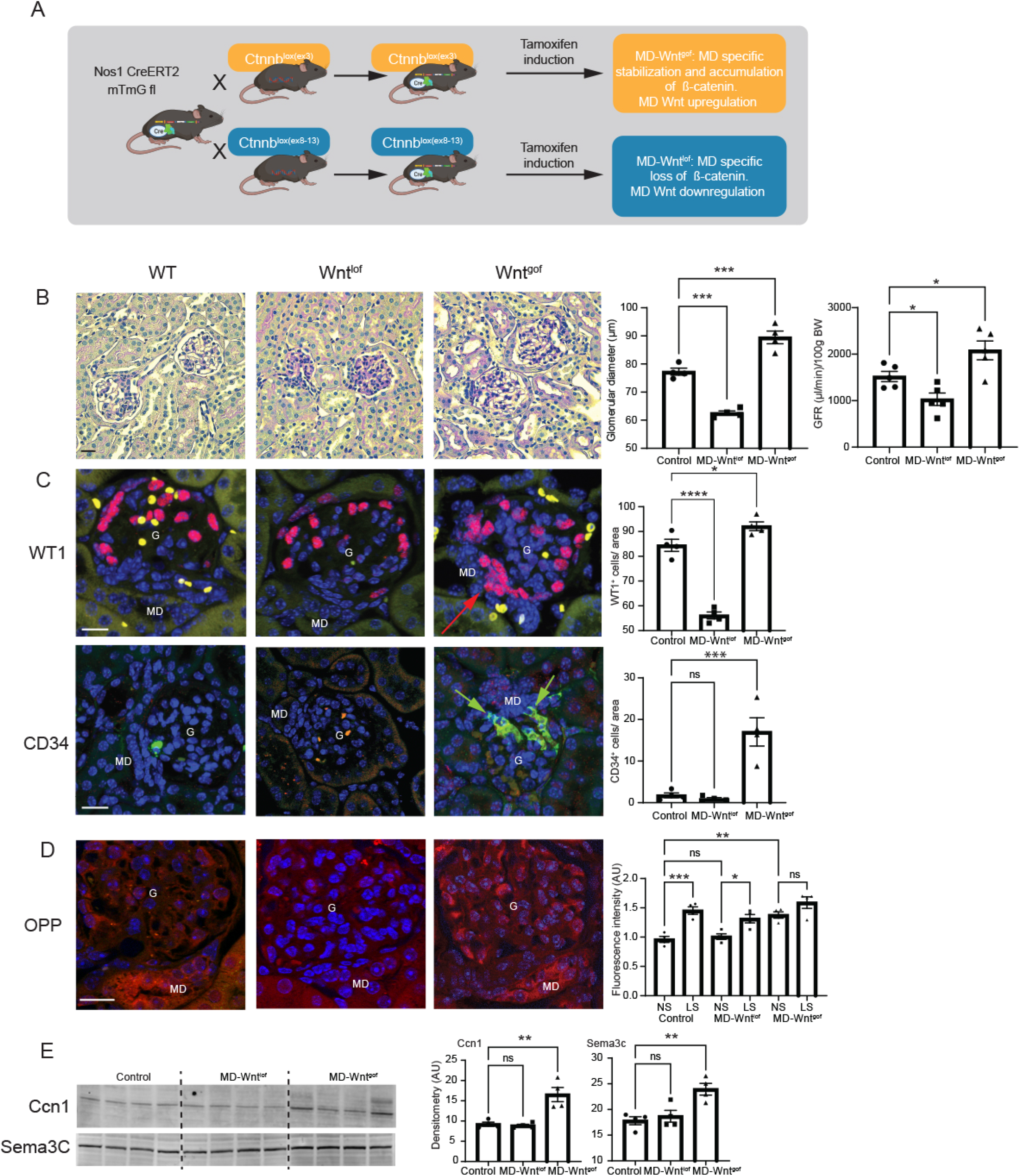
The effects of genetic manipulation of MD cell Wnt signaling on glomerular structure and function. **(A)** Illustration of the applied Cre/lox-based breeding strategy to generate MD Wnt gain-of-function (MD-Wnt^gof^) and loss-of-function (MD-Wnt^lof^) mouse models. **(B)** Renal histological (representative H&E images, left) and functional (glomerular filtration rate (GFR), right) features of MD-Wnt^gof^ and ^lof^ mice 2 months after tamoxifen induction. Note the enlarged, hypercellular and smaller, hypocellular cortical glomeruli in MD-Wnt^gof^ and ^lof^ mice, respectively, compared to control WT. Glomerular capillary lumen appears reduced with extracellular (mesangial) matrix accumulation in MD-Wnt^lof^ mice. Bar is 25µm. **(C-D)** Representative immunofluorescence images (left) and statistical summary (right) of WT1^+^ (red) and CD34^+^ (green) cell number (C) and quantitation of protein synthesis (D, OPP staining, red). Note the high cell density at the macula densa (MD) cell base (arrows, C) and OPP intensity (red) in MD cells indicating high level of global protein synthesis. Bar is 25µm. G: glomerulus. NS: normal salt, LS: low salt diet. Nuclei are labeled blue with DAPI. **(E)** Altered expression of MD-specific proteins in renal cortical homogenates including Ccn1 and Sema3c. Ns: not significant, *: p<0.05-****p<0.0001, data are mean ± SEM, n=5 mice each group, histology data represent the average of n=5 glomeruli from n=4 mice each.

### Human translation

To study the MD tissue regenerative program in the human kidney and its alterations in chronic kidney disease (CKD), we first analyzed the expression of the angiogenic factor CCN1 in freshly nephrectomized and fixed human kidney tissues from patients with normal kidney function or CKD (eGFR (estimated glomerular filtration rate) <50 mL/min/1.73m^2^). CCN1 protein expression was entirely specific to MD cells in the normal human renal cortex and medulla (Fig. 4A). Importantly, the number of CCN1^+^ MD cells was significantly reduced to almost undetectable levels in CKD (Fig. 4A).

**Figure 4.**
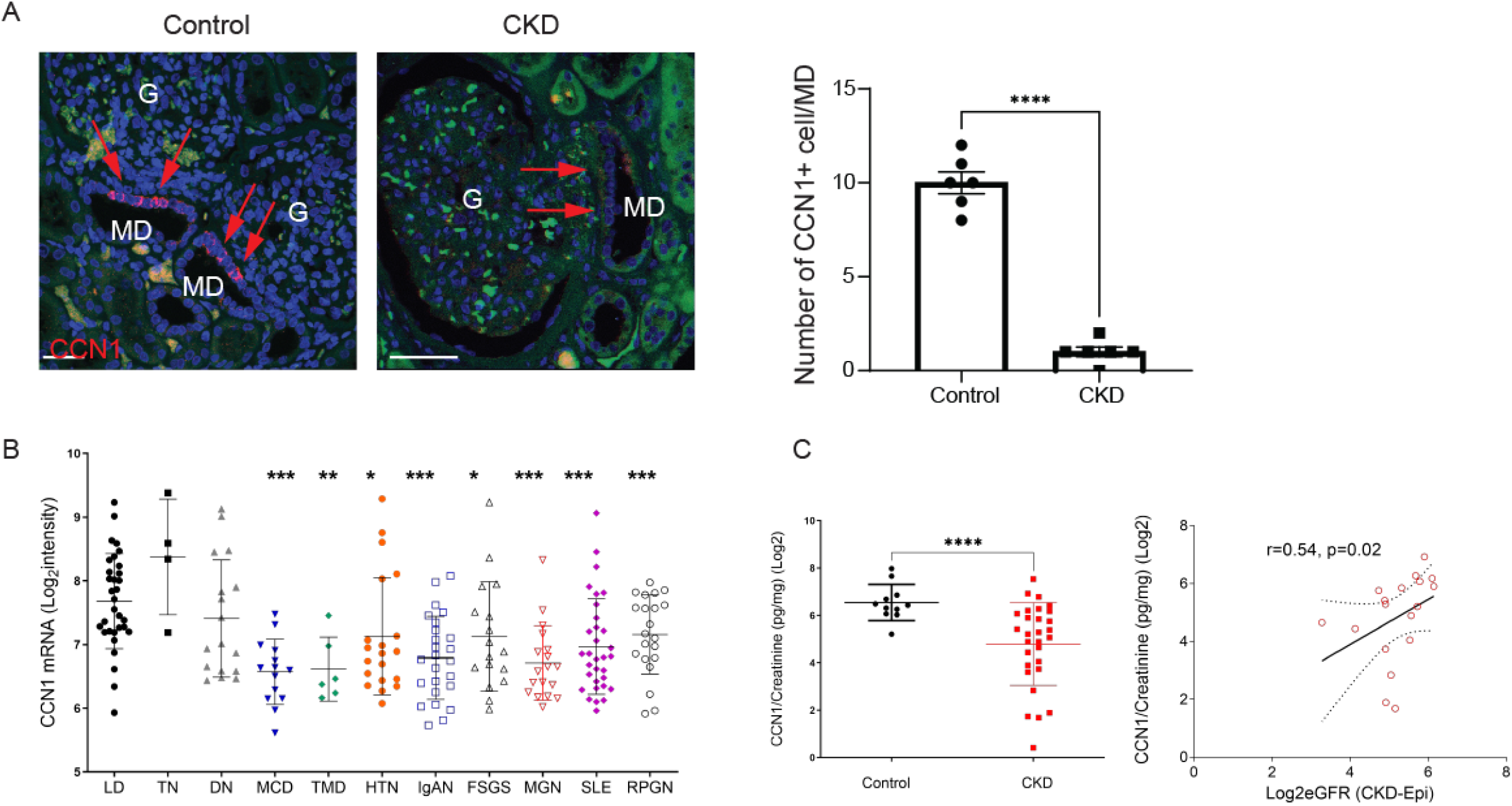
Human kidney expression of CCN1 in patients with normal kidney function or CKD. **(A)** Immunofluorescence labeling (red, left) and quantification (right) show strong CCN1 expression exclusively in cells of the macula densa (MD, red arrows) in control, while labeling is mostly absent in kidney tissue samples from patients with CKD. Nuclei are labeled blue with DAPI. G: glomerulus. Bar is 25µm. *: p<0.05, data are mean ± SEM, each data point represents the average of n=10 glomeruli per kidney, n=6 patients each. **(B)** Intrarenal CCN1 (CYR61) mRNA expression in kidney biopsies (tubulointerstitial compartment) from living donor (LD), tumor nephrectomy (TN), and CKD patients with various etiologies from the European Renal cDNA biobank (ERCB). LD (n=31), TN (n=4), Diabetic Nephropathy (DN, n=17), Minimal Change Disease (MCD, n=14), Thin Basement Membrane Disease (TMD, n=6), Arterial Hypertension (HTN, n=20), IgA Nephropathy (IgAN, n=25), Focal Segmental Glomerulosclerosis (FSGS, n=17), Lupus Nephritis (SLE, n=32), Membranous Glomerulonephropathy (MGN, n=18), Vasculitis (RPGN, n=21). Differential expression comparison between LD and each disease subtype was performed using T-test, data are mean ± SEM, *, ** and *** denotes p<0.05, <0.01, and p<0.001, respectively. **(C)** The association of urinary CCN1 levels with kidney function. Comparison of urinary CCN1 in control (n=11) and CKD patients (n=29, p < 0.0001) (left) and the positive correlation between urinary CCN1 excretion and eGFR in CKD patients (n=18, log2 transformed urinary CCN1/creatinine ratios are shown)(right).

Transcriptomic data in human kidney biopsies from the European Renal cDNA Biobank (ERCB) was analyzed to further investigate the relevance of CCN1 to human CKD. The results indicated that compared to living donors, CCN1 was significantly under-expressed in CKD patients with minimal change disease, thin basement membrane, membranous glomerulonephropathy, IgA nephropathy, lupus nephritis, vasculitis, hypertensive nephropathy, and focal segmental glomerulosclerosis (Fig. 4B). In fact, the results placed CCN1 within the top 1% of under-expressed genes in CKD.

In addition, we tested whether CCN1 can be detected in human urine samples and if urinary CCN1 levels correlate with kidney function, thus a potential biomarker. The results confirmed the presence of detectable levels of urinary CCN1 excretion, and importantly, urinary CCN1 was significantly diminished in CKD patients (4.8 ± 0.3, log2 transformed CCN1/Creatinine ratio) compared to volunteer controls (6.5 ± 0.2) (Fig. 4C). Furthermore, urinary CCN1/Creatinine levels showed a significant positive correlation with eGFR (Fig. 4C) suggesting that reduced urinary levels of CCN1 are associated with loss of kidney function. In addition, patients with lower CCN1/Creatinine levels (defined as less than average value among this group of patients) showed a trend to have lower eGFR, older age, and were more likely to progress to end stage kidney disease (not shown).

### Therapeutic targeting of the MD program for kidney repair

To study whether targeting of the presently identified MD cell program for kidney tissue regeneration provides therapeutic benefit, the effects of treatment with MD-derived biologicals were tested in the robust CKD model of Adriamycin (ADR)-induced glomerulosclerosis (GS) in BalbC mice (Fig. 5A). A single ADR injection induced severe GS pathology after two weeks indicated by reduced GFR and high-level albuminuria (Fig. 5B-C). At this point, treatment of the pre-existing GS was initiated using one of the following five biologicals: saline (PBS), recombinant CCN1 in low or high-dose, control DMEMF12 culture medium or conditioned cell culture medium of the recently developed and characterized MD^geo^ MD cell line (*7*)(Fig. 5A). Animals were followed-up for 4 weeks of treatment. Mass spectrometry analysis and CCN1 ELISA of the conditioned MD^geo^ cell culture medium confirmed the secretion of MD cell factors and informed the therapeutic dose of low CCN1 (Fig. S3A), while the high-dose CCN1 was chosen based on previous work in liver (*14*). In contrast to control PBS or DMEMF12 medium which had no effect, treatment with CCN1 (either with low or high dose) or conditioned MD medium caused strong improvements in albuminuria (Fig. 5C). In contrast to all other groups, treatment with MD medium dramatically improved GFR (Fig. 5B). Subsequent kidney histological analysis showed severe GS and tissue fibrosis in control PBS and DMEMF12-treated groups, while CCN1 or MD medium treatments greatly improved GS pathology, p57^+^ podocyte number (Fig. 5D-F) and tubulointerstitial fibrosis (Fig. S3B-C).

**Figure 5.**
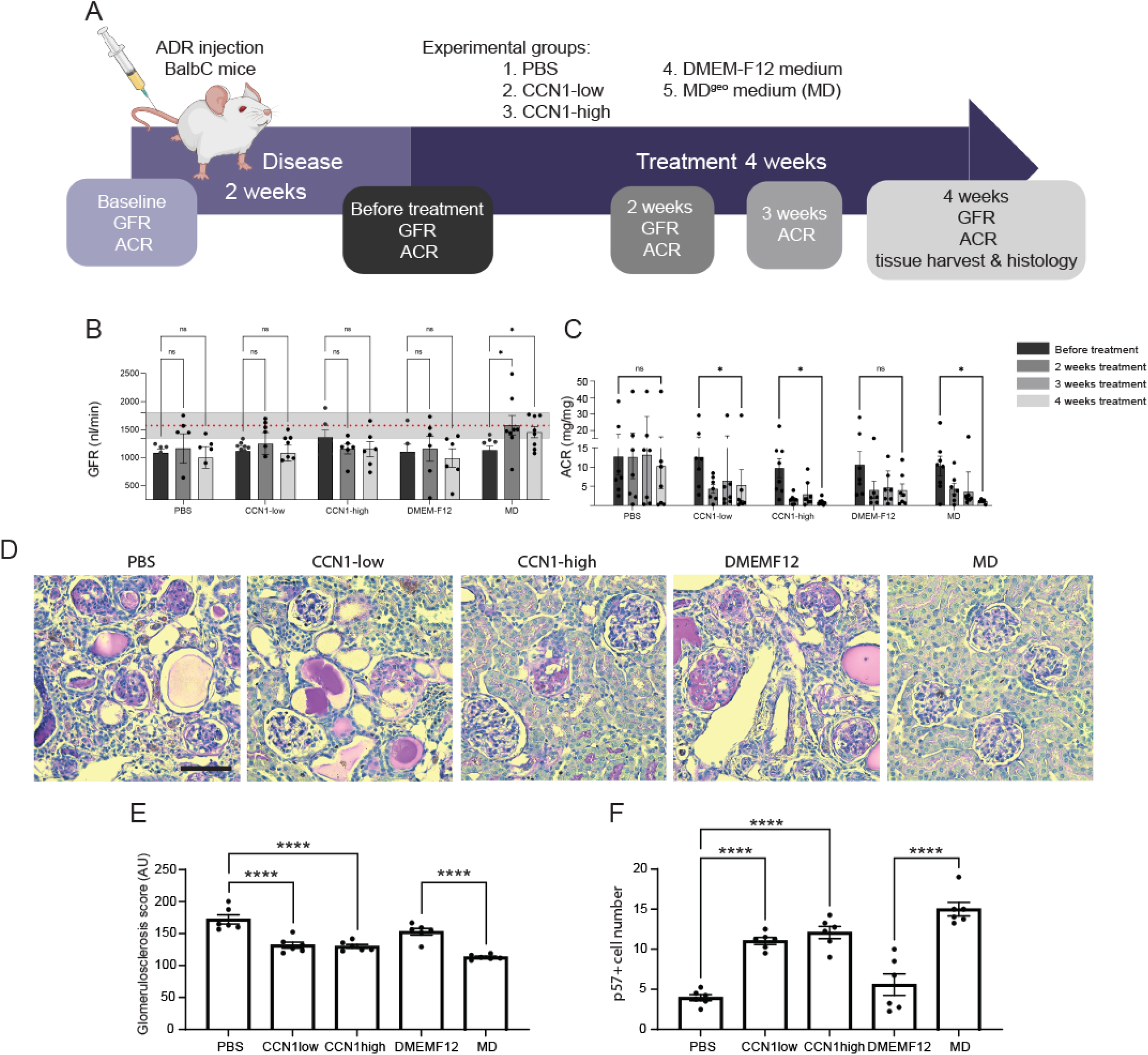
Dramatic improvement of pre-existing CKD in response to treatment with MD biologicals. **(A)** Illustration of therapeutic study design for testing the effects of MD biologicals using the Adriamycin (ADR) model of glomerulosclerosis in BalbC mice. **(B)** Time-course of GFR changes followed in the same mice measured by the MediBeacon non-invasive transcutaneous method. Note the significant improvement of GFR returning to normal baseline levels (red dotted line representing mean ± SEM (grey shaded area), measured at baseline) in the MD treatment group indicating functional regression of FSGS pathology. **(C)** Time-course of albuminuria (Albumin/Creatinine Ratio, ACR) changes followed in the same mice measured by ELISA. Note the significant improvement in albuminuria in the CCN1 and MD treatment groups in contrast to the PBS and DMEM-F12 controls. **(D)** Representative images of PAS-stained histological sections of kidney tissues harvested at the end of treatment in all treatment groups. **(E)** Quantification of glomerulosclerosis evaluated blindly by measuring the density of PAS staining on histological sections using Image J. **(F)** Podocyte preservation calculated in tissue volumes based on the number of p57^+^ cells. Ns=not significant, *P<0.05, **P<0.01, ***P<0.001, ****P<0.0001, n=6-8 mice each, histological data points represent the average of n=20 glomeruli per animal, n=6 mice each.

## Discussion

The newly discovered role of MD cells in endogenous kidney tissue remodeling and repair is a significant advance in the renal and cardiovascular field and promises the development of new regenerative therapeutic approaches for kidney and potentially other organ diseases. The concept that organ-specific physiological activation of chief cells orchestrate resident progenitor cell-mediated endogenous tissue repair has strong foundations in evolutionary biology, physiology, and clinical nephrology. The loss and conservation of body fluid and salt is a known major driver of the evolution of the mammalian nephron (*6*), major physiological adaptations to maintain body fluid homeostasis and blood pressure (*15*), and in the protective effect of low dietary salt intake (and ACEi) to slow down CKD progression (*16, 17*). Some of the presently identified MD mechanisms (e.g. Wnt signaling and CCN1) have well-known roles in the remodeling and repair of many tissues and organs (*12, 18*), while others (Aard, Pamr1) are yet to be explored in the kidney and beyond.

MD cells are chief salt sensors and regulators of kidney function, but our knowledge of these unique cells has been limited to their role in maintaining renal hemodynamics and activity of the renin-angiotensin system (*15*). The distinctively high MD cell MAPK (*19*), mTOR (*20*), calcium (*7*), and now Wnt signaling activity (Figs. 3 and S2A-B) among all renal cell types, and their recently identified neuron-like features (*7*) underscore that MD cells function as the nephron central command. However, their major tissue regenerative function, detailed gene profile and the molecular identity of MD-derived secreted tissue remodeling factors have been unknown. The reasons for this include the inaccessible, minority cell type that has been difficult to study, technology limitations in the past, and likely the elements in the western diet (e.g. high dietary salt intake) that may blunt this endogenous regenerative mechanism.

Our group recently established serial intravital MPM imaging to track the fate of individual, genetically labeled renal cell types over several days in the intact kidney (*8-10, 21*), which technique was applied in the present work for different mesenchymal and endothelial precursor cell types and in renal injury (Fig. S1A) and physiological activation conditions (Fig. 1A). This technological advance and approach provided critically valuable visual clues regarding the dynamic (within two weeks) and the MD-centric pattern of the recruitment and clonal propagation of vascular pole progenitor cells (Figs. 1A and S1C). Complemented with histological analysis and the glomerular vascular pole accumulation of WT1^+^ and CD34^+^ cells (Fig. 3B) that are known mesenchymal /podocyte and endothelial progenitor cell markers (*22, 23*) under MD stimulation conditions, these results strongly suggested the MD secretion and paracrine actions of local tissue remodeling factors that control the activation, recruitment and differentiation of resident progenitor cells. Vascular pole endothelial progenitors and cells of the renin lineage have been recently identified as local progenitor cells for glomerular cell types (*10, 24*). The appearance and clonal expansion of a WT1^+^/Renin^+^ mesenchymal cell population under the MD base in response to LS+ACEi or MD-Wnt^gof^ (Figs. S1C and 3B) are consistent with the important role of MD cells, specifically MD Wnt signaling, and with the recently established role of WT1 in the proliferation, migration, and differentiation of the cells of the renin lineage toward podocyte fate (*21*). The MD point-source origin of tissue remodeling factors may provide for a 3D structured tissue remodeling program in each nephron that likely involves a complex balancing act between inflammatory/pro-fibrotic and regenerative/anti-fibrotic mechanisms. The known biological activities of the presently identified multiple secreted MD tissue remodeling factors (Fig. 2) is supportive of this view. The MD-centric progenitor cell recruitment and tissue remodeling pattern suggests the involvement of MD-derived mechanism in the previously described progenitor/podocyte cell marker gradient (*25*) and podocyte precursor cell accumulation in the vascular pole region of Bowman’s capsule including at young age (*26, 27*) and in response to ACEi treatment (*28*). Cox-2 and Nos1 dependence of mesenchymal and endothelial progenitor cell recruitment (Fig. 1B) is consistent with the role of MD cell factors, including PGE2 that was shown to activate the recruitment and differentiation of CD44^+^ mesenchymal progenitors to the renin cell fate (*29*).

The other technology advance that provided critically important new information on MD cell function in the present work is their detailed high-resolution transcriptome analysis using both bulk and single-cell RNA sequencing (Fig. 2). Until now, only a few studies reported limited MD transcriptome data using a very low number of cells (*30-32*). The high MD expression of secreted angiogenic, cell migration and patterning, growth, and ECM remodeling factors (Fig. 2B-D) is a molecular-level confirmation of the intravital progenitor cell-tracking results (Figs. 1 and S1) and provides further strong support for the newly identified tissue regenerative role of MD cells. Consistent with the activation of the MD tissue remodeling program by physiological stimuli (LS+ACEi treatment), the same stimulation increased the expression of key MD-specific factors underlying this function (Fig. 2D, Supplement Table 1). The top MD-enriched angiogenic (*Pappa2, Pamr1, Sfrp1, Vash2, Vegfa, Ccn1*), cell migration and patterning (*Unc5d, Sema3c, Robo2, Slit2*), growth (*Bmp3, Egf, Fgf9*) and ECM signaling and remodeling (*Frem1, Thsd4, Mmp14, Adamtsl2*) factors and gene networks, cytokines and chemokines have well-defined roles in angiogenesis, progenitor cell recruitment and differentiation, extracellular matrix and tissue remodeling, and regeneration of several organs (*18, 33, 34*). The MD secretion of several of these factors including the anti-fibrotic angiogenesis modulator Ccn1 was confirmed in the present study (Fig. S3A). While the secretion of these tissue remodeling factors by MD cells is in general consistent with the observed MD-centric pattern of mesenchymal and endothelial progenitor cell recruitment (Figs. 1 and S1C), identification of their specific function and contribution to the MD tissue remodeling and repair program requires future work. The presence of a dense network of long, basal MD cell processes that make contact with various target cells at the glomerular vascular pole and contain secretory cargo including Pappa2 (*11*) is consistent with the presently identified new MD cell function. In addition, our recent study demonstrated that among all renal cell types, MD cells have the highest rate of protein synthesis that is further stimulated by low salt diet (*20*), suggesting that MD cells synthesize and secrete the above factors under both control and stimulated conditions. The increased synthesis of global and specific MD proteins in MD-Wnt^gof^ mice (Fig. 3D-E) further supports this MD function.

Both pharmacological (Cox-2 and Nos1 inhibitors, Fig. 1B) and genetic approaches (MD-Wnt^gof/lof^ mice, Fig. 3) validated the MD-specificity of the newly discovered endogenous tissue remodeling pathway. While the role of Wnt in this new MD mechanism was much expected due to the high expression of several Wnt target genes (Fig. 2C-D) and its well-known function in tissue regeneration (*12*), additional major signaling pathways are likely involved. Several molecular players of Notch, Shh, FGF, Hippo signaling are represented in the top enriched genes in the MD transcriptome (Fig. 2C-D, Supplement Table 1). Future work is needed to dissect their roles in MD cells. The known structural and neuron-like functional heterogeneity between individual MD cells (*7, 11, 35*) was reflected also in their regenerative gene profile established by single-cell transcriptome analysis (Fig. 2C). The clustering of angiogenic, ECM remodeling, precursor, etc. MD cell subtypes (Fig. 2C) suggests that there may be individual MD cells dedicated to controlling endothelial, mesenchymal progenitor cells or self-renewal separately. However, long-term genetic fate tracking of MD cells using inducible MD-GFP reporter mice did not find labeled cells outside the MD (data not shown), that suggests lack of MD cell differentiation to other renal cell types, at least in the mouse kidney.

The newly acquired knowledge on MD cells and their new functions suggest that these cells may play an important role in human kidney disease, and that targeting and augmenting MD mechanisms may improve CKD pathology. Therefore, additional studies were performed to translate the new MD function to the context of human kidney disease (Fig. 4) and therapeutic interventions (Fig. 5). CCN1 was specifically expressed in MD cells in the human kidney (Fig. 4A), and its expression was reduced in non-diabetic CKD based on both mRNA (Fig. 4B) and protein levels (Fig. 4A). CCN1 was detectable in human urine samples and correlated with glomerular function (Fig. 4C), suggesting potential utility and need for further development of MD-derived urinary CCN1 as a new non-invasive biomarker of glomerular structure and function and CKD progression.

The availability and use of the recently established new MD cell line MD^geo^ (*7*) was yet another critically important tool in this study that allowed the testing of the therapeutic potential of MD-derived biologicals (Fig. 5). The use of either a specific, single (Ccn1) or a mixture of all MD factors (conditioned medium) provided initial proof-of-concept of the potent protective effects of targeting MD cell mechanisms. MD-derived biologicals are readily available from the conditioned MD^geo^ culture medium (Fig. S3A) that provided ease of therapeutic use in addition to recombinant proteins (Ccn1). Compared to before treatment, the dramatically improved albuminuria, GFR, and renal histopathology in response to treatment with conditioned MD medium (and partially Ccn1) suggests functional and structural regression of pre-existing CKD (Figs. 5B-F, S3B-C). The dramatic improvements in GS and interstitial fibrosis observed in animals treated with CCN1 or MD medium (Figs. 5D-F, S3B-C) is consistent with the known anti-fibrotic function of CCN1 (*14, 18*) and other MD-derived factors. It should be emphasized that the presently applied MD stimuli (LS+ACEi treatment, inducible MD-Wnt^gof^, MD biologicals) that resulted in improved kidney structure and function were all applied temporarily. However, sustained overactivation of the MD including the presently identified endogenous tissue regenerative mechanism may become pathogenic as seen for example in early diabetes (*36*). To determine whether this new MD mechanism is involved in the protective effects of SGLT2 inhibitors (*36*), or if its stimulation or inhibition is relevant to different phases of diabetic or non-diabetic kidney disease requires further study. It is likely that the ultimate protective or pathogenic role, and therefore the potential therapeutic targeting of the new MD tissue regenerative program is context dependent.

In summary, the present study uncovered a novel endogenous tissue remodeling mechanism in the kidney that is activated by the physiological stimulation of the chief MD cells that function as the nephron central command. The MD program for kidney tissue remodeling and repair controls resident progenitor cells via secreted angiogenic, growth and extracellular matrix remodeling factors, cytokines and chemokines. The identification and molecular characterization of this new MD mechanism provided here represents a key step in developing and targeting this pathway in the future for potential diagnostic and therapeutic purposes.

## Materials and Methods

### Animals

Male and female, 6-12 weeks old C57BL6/J or BalbC mice (Jackson Laboratory, Bar Harbor, ME) were used in all experiments. Transgenic mouse models with the expression of various fluorescent reporter proteins and gene knock-out strategies were generated by intercrossing Cre or Cre-ER^T2^ mice with flox mice as indicated in the table below.

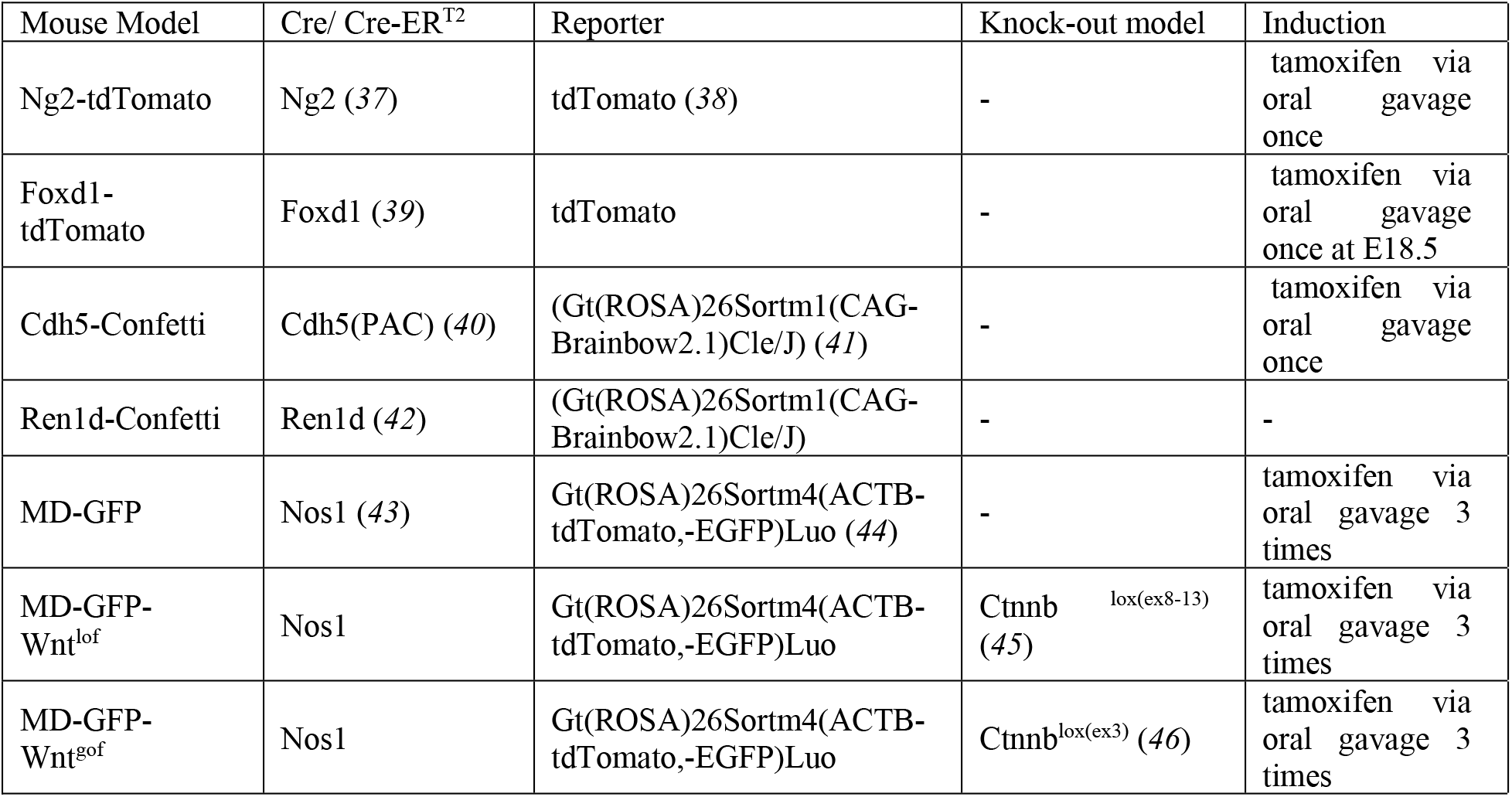
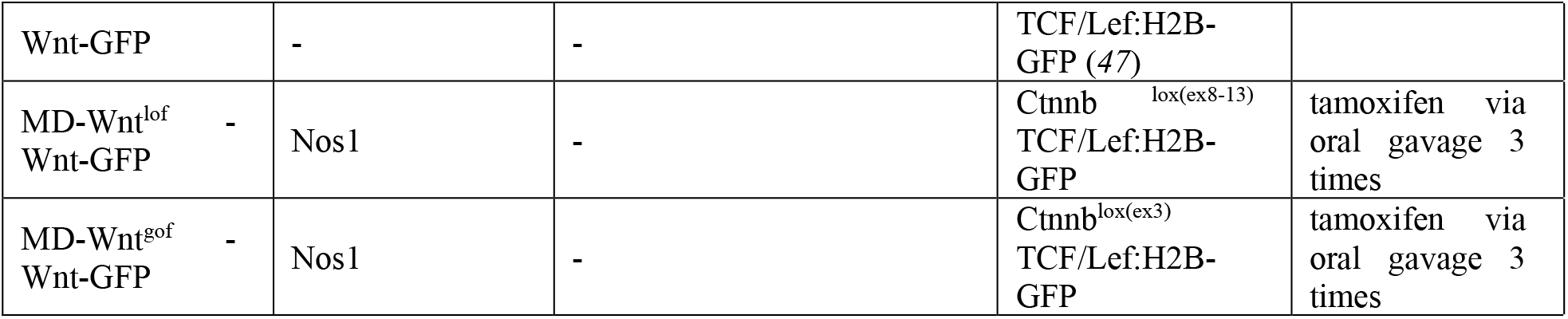

Tamoxifen was administered by oral gavage (75mg/kg) once or a total of three times (every other day) in the Cre-ER^T2^ mouse models resulting in cell specific expression of reporters and gene knock-out mouse models. Some mice received low-salt diet for 10 to 14 days (TD 90228, Harlan Teklad, Madison, WI) with or without ACE inhibitor treatment using enalapril (150 mg/L via drinking water, E6888 purchased from Sigma Aldrich Solutions, MO) with or without selective COX-2 inhibitor (SC58236, 6mg/L via drinking water, from Pfizer Inc. New York, NY), or selective Nos1 inhibitor 7-NI (20mg/kg ip daily injections, N778, Sigma Aldrich, St. Louis, MO); or furosemide (30mg/kg via ip injections daily, Chinoin, Budapest, Hungary). None of the treatments resulted in significant alterations in blood pressure based on tail-cuff measurements. All animal protocols were approved by the Institutional Animal Care and Use Committee at the University of Southern California.

### Unilateral IRI

Unilateral clamping of the left renal pedicle was performed in anesthetized mice using Isoflurane (1-2% inhalant via nosecone) and body temperature was controlled at 37°C during surgery with a temperature-controlled operating table (SomnoSuit Low-Flow Anesthesia System, Kent Scientific, Torrington, CT). The left kidney was exteriorized through a 0.5 cm long flank incision and the left renal pedicle was dissected and clamped for 30 minutes of ischemia followed by the release of the clamp. The left kidney was inserted back into the retroperitoneum, and the wound was sutured. Animals that underwent only left kidney exteriorization but not left renal pedicle clamping (Sham operation) served as control. At the end of 2 weeks follow up, mice were euthanized, and tissues harvested for histological analysis.

### Adriamycin nephropathy

To establish the animal model of Adriamycin (ADR) induced nephropathy, Balb/C mice at 8 weeks of age received a single iv injection of ADR (10.5 mg/kg, Sigma Aldrich, St. Louis, MO) via tail vein injection and were followed for a total of 6 weeks (*48*). At the 2 weeks peak of disease when glomerulosclerosis pathology and albuminuria were stabilized, 4-weeks treatment of pre-existing pathology was initiated in the following 5 groups by daily ip injections: (i) control PBS, (ii) low-dose CCN1 (angiogenic modulator, 5 ng/day), (iii) high-dose CCN1 (1 mg/kg/day), control DMEM-F12 cell culture medium, and (v) conditioned macula densa (MD) cell culture medium. At the end of 4 weeks treatment, mice were euthanized, and tissues harvested for histological analysis.

### Serial intravital imaging using multiphoton microscopy (MPM)

Surgical implantation of a dorsal abdominal imaging window (AIW) above the left kidney was performed on *NG2-tdTomato* and Cdh5-Confetti mice using aseptic surgical procedures as described recently (*49*). This approach allows the non-invasive imaging of the same glomeruli over time for up to 30 days (*24*). For repeated MPM imaging of the same mice with AIW, animals underwent brief anesthesia sessions every 3–4 days using 1%–2% isoflurane and the SomnoSuite low-flow anesthesia system (Kent Scientific, Torrington, CT). Body temperature was maintained with a homeothermic blanket system (Harvard Apparatus). Alexa Fluor 594 or 680-conjugated albumin (Thermo Fisher, Waltham, MA) was administered iv. by retro-orbital injections to label the circulating plasma (30 µL iv. bolus from 10 µg/ml stock solution). The images were acquired using a Leica SP8 DIVE multiphoton confocal fluorescence imaging system with a 63× Leica glycerine-immersion objective (numerical aperture (NA) 1.3) powered by a Chameleon Discovery laser at either 860 nm or 970 nm (Coherent, Santa Clara, CA) and a DMI8 inverted microscope’s external Leica 4Tune spectral hybrid detectors (emission at 460-480nm for CFP, 510-520 nm for eGFP, 550-570 nm for YFP, 580-600 nm for tdTomato, 600-620 nm for RFP) (Leica Microsystems, Heidelberg, Germany). The potential toxicity of laser excitation and fluorescence to the cells was minimized by using a low laser power and high scan speeds to keep total laser exposure as minimal as possible. The usual image acquisition (12-bit, 512×512 pixel) consisted of only one z stack per tissue volume (<2 min), which resulted in no apparent cell injury. Fluorescence images were collected in volume series (xyz, 1 s per frame) with the Leica LAS X imaging software and using the same instrument settings (laser power, offset, gain of all detector channels). Serial imaging of the same glomerulus in the same animal/kidney was performed once every 3 to 4 days for up 14 days after the first imaging session. Maximal projections from *Z*-stacks were used to count and compare tdTomato and Confetti^+^ cell number in the same area over time. In the Confetti mouse models clonal or monochromatic tracing units were defined as numerous directly adjacent individual cells that featured the same Confetti color combination. All ten possible Confetti color combinations were observed as described before (*10*). The counting of tdTomato or Confetti^+^ cells and clones was facilitated by standardized image thresholding using ImageJ (NIH), Leica LAS X (Leica Microsystems Inc.), and cell-counting algorithms of Imaris 9.2 3D image visualization and analysis software (Bitplane USA) for intravital imaging *Z*-stacks.

### Global protein synthesis assay using o-propargylpuromycin fluorescence imaging and quantification

Global protein synthesis was assessed by O-propargyl-puromycin (OPP) labeling (Thermo Fisher Scientific) as described before (*20*). Briefly, mice were injected intraperitoneally with 25 uL of 20 mM OPP 1 h before tissue harvest and staining. For OPP labeling, sections were developed using the Alexa Fluor 594 Click-iT OPP Protein Synthesis kit (Thermo Fisher Scientific) according to the manufacturer’s instructions. OPP fluorescence intensity was quantified by imaging all tissue sections using identical laser power and confocal microscopy settings. Fluorescence intensity was measured by placing 10 regions of interest across the entire MD plaque (FOPP) and normalized to the fluorescence intensity of red blood cells (RBCs) (FRBC). The average FOPP of 5–10 different representative MD cell plaques was quantified in each tissue section.

### Glomerular filtration rate measurement

GFR measurements were performed using the MediBeacon Transdermal Mini GFR Measurement System (MediBeacon) as described previously (*50*). Briefly, mice were anesthetized, MediBeacon sensor was placed on the depilated dorsal skin. Mice were injected retro-orbitally with the inulin analog exogenous GFR tracer fluorescein-isothiocyanate conjugated sinistrin (FITC-S 7.5 mg/100 g body weight, MediBeacon, St. Louis, MO). The excretion kinetics of the FITC-S was measured for 90 minutes. GFR was then calculated based on the decay kinetics (half-life time) of FITC-S using MediBeacon Data Studio software (MediBeacon).

### Blood pressure measurement

Systolic blood pressure was measured by tail-cuff plethysmography (Visitech BP-2000, Visitech System Inc.) in trained animals as previously described (*19*).

### Tissue processing, immunofluorescence, and histology

Immunofluorescence detection of proteins was performed as described previously (*19*). Briefly, cryosections were cut at 25 µm, washed with 1x PBS. Paraffin tissue blocks were sectioned to 8 µm thick. For antigen retrieval, heat-induced epitope retrieval with Sodium Citrate buffer (pH 6.0) or Tris-EDTA (pH 9.0) was applied. To reduce non-specific binding, sections were blocked with normal serum (1:20). Primary and secondary antibodies were applied sequentially overnight at 4° C and 2 hours at room temperature. Primary antibodies and dilutions were as follows, anti-villin antibody (1:100, sc58897, Santa Cruz Biotechnology, Dallas, TX), anti-claudin1 antibody (1:100, SAB4503546, Sigma Aldrich, St. Louis, MO), anti-renin antibody (1:100, AF4277, R&D Systems, Minneapolis, MN), anti-alpha smooth muscle actin antibody (αSMA, 1:100, A2547, Sigma Aldrich, St. Louis, MO), anti-platelet-derived growth factor receptor beta antibody (PDGFRβ, 1:100, 3169S, Cell Signaling Technology, Danvers, MA), anti-podocin antibody (1:100, SC22298, Santa Cruz Biotechnology, Dallas, TX), anti-CCN1 antibody (1:100, SC13100, Santa Cruz Biotechnology, Dallas, TX), anti-growth differentiation factor 15 antibody (GDF15, 1:100, HPA011191, Sigma Aldrich, St. Louis, MO), anti-pappalysin2 antibody (PAPPA2, 1:100, PA5-21046, Invitrogen, Waltham, MA), anti-Wilms tumor protein antibody (WT1, 1:100, AB89901, Abcam, Cambridge, UK), anti-CD34 antibody, (1:100, AB8536, Abcam, Cambridge, UK), anti-active β-catenin antibody (ABC, 1:100, 05-665, Millipore, Burlington, MA), anti-Meis2 antibody (1:100, HPA003256, Sigma Aldrich, St. Louis, MO). Alexa Fluor 488, 594, and 647-conjugated secondary antibodies were purchased from Invitrogen. Slides were mounted by using DAPI-containing mounting media (VectaShield, Vector Laboratories Inc., Burlingame, CA). Vectastain Elite ABC-HRP Kit was used to perform horseradish-peroxidase labeling (PK6101, Vector Laboratories, Burlingame, CA). Sections were examined with Leica TCS SP8 (Leica Microsystems, Wetzlar, Germany) confocal/multiphoton laser scanning microscope systems as described previously (*11*). Imaging software (Image J, National Institutes of Health and the Laboratory for Optical and Computational Instrumentation (LOCI, University of Wisconsin) was used to calculate the percent area of colocalization to determine the changes in the proportion of NG2^+^ cells in each cell type, as well as the proportional distribution of NG2^+^cell types.

For the assessment of tubulointerstitial fibrosis, histological analysis of Picrosirius red (PSR) staining was performed on mouse kidney sections using Sirius Red F3B (Sigma-Aldrich, St. Louis, MS). For the evaluation of glomerulosclerosis, histological analysis of periodic acid-Schiff stain was performed on mouse kidney sections using PAS Stain Kit (24200-1, Polysciences, Warrington, PA). Images were visualized at 25× magnification using Leica TCS SP8 (Leica Microsystems, Wetzlar, Germany). Imaging software (Image J) was used to calculate the percent area of red-stained collagen or PAS positive areas as described before (*51*).

### Tissue CLARITY

Three-dimensional imaging was performed as previously described (*52*) by carrying out whole-mount immunofluorescence stains on slices of MD-GFP WT, Wnt^gof^, and Wnt^lof^ mouse kidneys. Slices were fixed in 4% formaldehyde in 1x phosphate buffer saline (PBS) at room temperature for 45 min, washed in 1XPBS, blocked in 1xPBS with 0.1% TritonX100 and 2% SEA Block (ThermoFisher Scientific) for 1 hour, and sequentially incubated in primary and secondary antibodies over 2 days. Primary antibodies and dilutions were as follows: anti-Wilms tumor protein antibody (WT1, 1:100, AB89901, Abcam, Cambridge, UK), anti-Meis2 antibody (1:100, HPA003256, Sigma Aldrich, St. Louis, MO). Primary and secondary antibodies were diluted in the blocking solution. To clear tissue slices, the slices were dehydrated in methanol via increasing concentrations 50%, 75%, 100%, diluted in PBS - each for 1hr - and subsequently submerged in a 50:50 benzyl benzoate/benzyl alcohol (BABB): methanol solution, followed by 100% BABB. High resolution imaging of MD plaques and the adjacent glomeruli was performed on a Leica SP8 multiphoton microscope using a 63X glycerol immersion objective.

### MD cell isolation

MD-GFP cells were isolated as described before (*11*). Briefly, kidney cortex was isolated from freshly harvested mouse kidneys and digested using Hyaluronidase and Liberase TM enzyme combination (concentration: 2mg/mL and 2.5mg/mL respectively, Sigma Aldrich, St. Louis, MO). After digestion, MD cells were isolated based on their genetic reporter expression (GFP) by using FACS ARIAII cell sorter, and excitation wavelengths 488 and 633nm in sterile conditions. The highest tdTomato expressing cells were collected as controls with high representation of distal tubule and collecting duct segments, podocytes, and vascular smooth muscle cells.

### RNA sequencing and bioinformatics

Whole-transcriptome RNAseq was performed at the USC Norris Molecular Genomics Core as described before (*7*). Cells were extracted using Qiagen miRNeasy purification kit following manufacturer’s protocol for total RNA purification (Qiagen cat#217004). Libraries were simultaneously prepared using Takara’s SMARTer Stranded Total-RNA Pico v2 library preparation kit following manufacturer’s protocol (Takara cat#634412). Prepared libraries were sequenced on Illumina Nextseq500 at 2×75cycles.

RNA-seq data was analyzed using the RNA-seq workflow in Partek Flow software (V10.1.21. Partek Inc., St. Louis, MO, USA). Briefly, the raw sequencing reads were first trimmed based on the quality score (Phred QC>=20, min read length=25 nt) before mapped to mouse genome build mm10 using Star 2.61 (*53*) with default parameter settings and Gencode M21 mouse transcriptome annotation (*54*) as guidance. Gencode M21 was then used to quantify the aligned reads to genes using Partek E/M method. Finally, gene level read counts in all samples were normalized using Upper Quartile normalization (*55*) and subjected to differential expression analysis using Partek Gene Specific Analysis method (genes with fewer than 10 aligned reads in any sample among a data set were excluded). The differentially expressed gene (DEG) lists were generated using the cutoff of FDR<0.05 and fold changes greater than 2.0 either direction. Z-scores based on the normalized gene counts from the top MD cell enriched genes in 5 categories were used to generate the heatmap in GraphPad Prism 9.0.1 (San Diego, California). Pathway analysis, graphical summary, biological and disease functions of IPA were used to analyze transcriptome data.

Single cell RNA sequencing was prepared using 10x Genomics 3’ v3.1 (cat# 1000092) following manufacturer’s protocol as described before. Samples were parsed into single cells using 10x Genomics Chromium Controller and libraries were simultaneously prepared. Prepared single cell RNA sequencing libraries were sequenced on the Illumina Novaseq6000 platform at a read length of 28×90 and read depth of 100,000 reads/cell for 2000-4000 cells. scRNAseq data was analyzed using the scRNAseq workflow by Partek Flow. Briefly, the raw sequencing reads with adaptors trimmed were mapped mm10 genome using Star 2.6.1 and quantified using Gencode M25 annotation to generate gene level counts. The gene counts were subjected to QA/QC and the low-quality cells were filtered using the following criteria: (1) contained less than 300 or more than 8000 detected genes, (2) mitochondrial counts higher than 15% of total counts. The counts were normalized using the Partek Flow recommended method (divided by 1 million, Add:1 and log2). Dimension reduction was carried out using PCA, followed by Graph-based Clustering with default settings and UMAP visualization (*56*). Cell populations were determined by expression of relevant biomarkers. A 5-fold thresholding in *Nos1* and *Pappa2* expression was applied to filter out potential non-MD cell contamination. *Nos1/Pappa2* expression filtered cells went through PCA, Graph-based clustering and UMAP visualization.

### Generation of conditioned MD^geo^ cell culture media

The recently established immortalized macula densa cell line (MD^geo^) was cultured as described before (*7*). Briefly, MD^geo^ cells were cultured in complete MD cell culture medium supplemented with nerve growth factor (NGF, 0.1 ug/mL; N8133, Millipore Sigma) at 33°C for proliferation. Cells were subcultured after 48 hours. For differentiation, cells were incubated at 37°C, 5% CO2 for 14 days in complete MD cell culture media supplemented with NGF (0.1ug/mL; N8133, Millipore Sigma). After full differentiation, MD^geo^ cells were physiologically activated by temporary exposure to low-salt conditions (*57*) for 6 hours every other day (3 times).

### Western Blot

For immunoblotting of mouse cortical homogenates, manually dissected slices of kidney cortex were homogenized in a buffer containing 20 mM Tris·HCl, 1 mM EGTA pH 7.0, and a protease inhibitor cocktail (BD Bioscience, San Jose, CA). Protein (40 µg) was processed for immunoblotting as described previously (*19*), using anti-CCN1 (1:500, SC13100, Santa Cruz Biotechnology, Dallas, TX), anti-Sema3C (1:500, MAB1728, R & D Systems, Minneapolis, MN) primary antibodies. After incubation, blots were incubated with secondary antibodies (1:5000; LI-COR Biosciences) and then visualized with Odyssey Infrared Imaging System (LI-COR Biosciences).

### Mass spectrometry analysis of the composition of MD^geo^ cell culture media

Cell-free MD^geo^ conditioned media were snap frozen and stored at −80°C. Proteins were precipitated using acetone. Proteins were digested and prepared using the SP3 protocol, with modifications, as previously described, using trypsin as a protease (*58*). Tryptic peptides were analyzed using a nanoscale liquid chromatography–tandem mass spectrometry hardware setup, consisting of a nanoflow LC (flow, 200 nl/min) coupled to an Orbitrap QExactive Plus tandem mass spectrometer. The peptides were separated using a gradient for reverse-phase separation, consisting of buffer A and buffer B, with ascending concentrations of buffer B (80% acetonitrile, 0.1% formic acid) over buffer A (0.1% formic acid). The peptides were separated using a 1-hour gradient.

Protein raw files were searched using MaxQuant and the LFQ algorithm (*59, 60*) with searches against a UniProt mouse proteome reference database released in January 2018. Search criteria were alkylation on cysteines as a fixed modification, and amino-terminal acetylation and methionine oxidation as variable modifications. Default criteria were used, meaning that PSM, peptide, and protein false discovery rates (FDRs) were set at 0.01. The LFQ algorithm was enabled, and “match between run” was enabled. The data were analyzed using Perseus version 1.5.5.3, with filtering for the embedded annotations as contaminant, reverse, or proteins identified by site only.

### CCN1 Assay

Mouse CYR61 ELISA Kit (ab253223, Abcam, Cambridge, UK) was used for the quantitative measurement of CCN1 protein in MD^geo^ cell culture media with or without conditioning according to the manufacturer’s instructions.

### Human studies

#### CCN1 protein analysis

Human MD cell CCN1 protein expression was quantified based on immunohistochemical analysis using anonymized adult (32-87 years old) formalin-fixed paraffin-embedded renal cortical tissues. Samples were obtained from unaffected regions of tumor nephrectomy specimens based on protocols HS-15-00298 and HS-16-00378 approved by the Institutional Review Board, Keck School of Medicine of the University of Southern California. Non-diabetic nephrectomy patients with either normal kidney function or CKD due to hypertensive nephropathy (eGFR <50 mL/min/1.73m^2^) were included. Basic patient history data regarding kidney function (eGFR) and comorbidities, such as hypertension, and medications were available.

The quantification method of CCN1 expression included the selection of 10 glomeruli in each sample in which the entire MD longitudinal section (glomerular mid-sections containing both vascular and urinary poles) was available. The number of CCN1^+^ individual MD cells per MD plaque were counted in a blinded fashion and the average of 10 MDs/glomeruli were used per kidney sample, n=6 patients (3 male and 3 female) each in control and CKD groups.

### Gene expression analysis

Transcriptomic data analysis of CCN1 was performed in human kidney biopsies from the European Renal cDNA Biobank (ERCB)(*61*). For tissue transcriptome analysis, gene expression profiles from 174 consecutive biopsies of patients with various CKD etiologies were compared with those from thirty-two living donor transplant biopsies obtained at the time of transplantation. Only patients aged 18 years or older were included in this study. All bio-specimens were procured after informed consent and with approval of the local ethics committees.

Transcriptome analysis was performed on microdissected tubulointerstitial components of human renal biopsies prospectively procured for molecular analysis, using Affymetrix GeneChip and TaqMan Low Density Arrays as previously published (*62*). Normalized expression data were log2-transformed and batch-corrected. FDR was applied to account for multiple testing.

### Urinary CCN1 analysis

Human CCN1 ELISA assay (R&D systems, Minneapolis, MN, USA) was used to analyze urinary CCN1 in undiluted samples collected from control (n=11) and patients with clinically confirmed CKD (n=29). Control samples were purchased from Bioreclamation IVT (New York, NY, USA). CKD samples were from the Clinical Phenotyping and Resource Biobank Core (C-PROBE) cohort based at University of Michigan that includes patients with CKD stage I-V. Spearman correlation analysis was performed between urinary CCN1/Creatinine levels and eGFR as described earlier (*63*). Molecular analysis of the renal biopsies, plasma and urine samples at University of Michigan Health System (UMHS) has been approved by the IRB of the University of Michigan (HUM00002468 and HUM00026609).

### Statistical methods

Data are expressed as average ± SEM and were analyzed using Student’s t-tests (between two groups), or ANOVA (for multiple groups) with post-hoc comparison by Bonferroni test. P<0.05 was considered significant. Statistical analyses were performed using GraphPad Prism 9.0c (GraphPad Software, Inc.).

## Acknowledgments

This work was supported in part by US National Institutes of Health grants DK064324, DK123564, and S10OD021833 to J.P-P. U.N.S. was funded by predoctoral research fellowship 19PRE34380886 of the American Heart Association. The bioinformatic work was partially supported by George M. O’Brien Michigan Kidney Translational Core Center, funded by NIH/NIDDK grant 2P30-DK-081943.

ERCB members at the time of the study included Clemens David Cohen, Holger Schmid, Michael Fischereder, Lutz Weber, Matthias Kretzler, Detlef Schlöndorff, Munich/Zurich/AnnArbor/New York; Jean Daniel. Sraer, Pierre Ronco, Paris; Maria Pia Rastaldi, Giuseppe D’Amico, Milano; Peter Doran, Hugh Brady, Dublin; Detlev Mönks, Christoph Wanner, Würzburg; Andrew Rees, Aberdeen and Vienna; Frank Strutz, Gerhard Anton Müller, Göttingen; Peter Mertens, Jürgen Floege, Aachen; Norbert Braun, Teut Risler, Tübingen; Loreto Gesualdo, Francesco Paolo Schena, Bari; Gunter Wolf, Jena; Rainer Oberbauer, Dontscho Kerjaschki, Vienna; Bernhard Banas, Bernhard Krämer, Regensburg; Moin Saleem, Bristol; Rudolf Wüthrich, Zurich; Walter Samtleben, Munich; Harm Peters, Hans-Hellmut Neumayer, Berlin; Mohamed Daha, Leiden; Katrin Ivens, Bernd Grabensee, Düsseldorf; Francisco Mampaso(†), Madrid; Jun Oh, Franz Schaefer, Martin Zeier, Hermann-Joseph Gröne, Heidelberg; Peter Gross, Dresden; Giancarlo Tonolo; Sassari; Vladimir Tesar, Prague; Harald Rupprecht, Bayreuth; Hermann Pavenstädt, Münster; Hans-Peter Marti, Bern; Peter Mertens, Magdeburg, Jens Gerth, Zwickau.

## Data availability

Data from bulk RNA-based MD transcriptome analysis is available at the Gene Expression Omnibus (GEO) repository at the National Center for Biotechnology Information (NCBI), with GEO accession number GSE163576. For single-cell RNA-based MD transcriptome analysis, GEO accession number is GSE189954.

## Author contributions

G.G. and J.P.P. designed the study, performed experiments, analyzed the imaging data, and wrote the manuscript. U.N.S, A.R.B, D.D., W.J., A.I., S.D., J.L.B., H.K., M.R., N.A., L.L., I.S.G., and M.K. performed experiments and made substantial contributions to acquire and analyze data. Y.C. filtered and analyzed transcriptome data. All authors approved the final version of the manuscript.

---

**Supplement Figure 1. Additional features of mesenchymal and endothelial precursor cell-mediated endogenous kidney tissue remodeling**.

**(A)** The effects of ischemia-reperfusion injury (IRI) or sham surgery in the ipsilateral or contralateral (cl) kidney (left) or treatment with furosemide, or low-salt (LS) diet or ACEi alone or in combination (LS+ACEi) (right) on Ng2^+^ cell number per glomerular area.

**(B)** Fate tracking of mesenchymal progenitor (Ng2^+^, red) cells in the renal cortex in timed control (top row) or after treatment with LS+ACEi for two weeks (center row). Representative fluorescence images of native Ng2^+^ cell tdTomato (red) with immunofluorescence co-localization of cell differentiation markers (green) for proximal tubule (villin), parietal epithelial (claudin-1), juxtaglomerular renin (renin), vascular smooth muscle (αSMA), and mesangial cells (PDGFRβ), and podocytes (podocin) in timed control or after LS+ACEi treatment (left). The relative distribution of the six differentiated cell types within the Ng2^+^ cell population is quantified in pie charts (right). The ratio of Ng2^+^ cells within each of the six differentiated cell types (% co-localization of double^+^/green cells) is shown below each cell type (bottom row). Note that the percentage of αSMA^+^ and PDGFRβ^+^ cells within the Ng2 lineage decreased, while all other cell types increased in response to LS+ACEi treatment.

**(C)** Representative fluorescence images of native tdTomato (red) in Foxd1^+^ cells (left) or Confetti (CFP/GFP/YFP/RFP multicolor) in Ren1d^+^ cells (right) in timed control or after LS+ACEi treatment in Foxd1-tdTomato or Ren1d-Confetti mouse kidney sections. Glomerular vascular pole areas under the MD cell base are indicated by arrows in multiple nephrons. Nuclei are labeled blue with DAPI. G: glomerulus. Scale bar is 50 μm for all panels.

**(D)** Quantification of Foxd1^+^ cell number and Ren1d^+^ or Cdh5^+^ clone frequency per glomerular area in timed control or after LS+ACEi treatment. All analysis was performed on frozen tissue sections. Ns: not significant, *: p<0.05-****p<0.0001, data are mean ± SEM, n=5 glomeruli averaged for n=4 mice each group.

**Supplement Figure 2. Validation of altered MD Wnt activity and its effects on systemic and renal parameters in MD-Wnt**^**gof**^ **and MD-Wnt**^**lof**^ **mice**.

**(A)** Representative images of activated β-catenin immunofluorescence (ABC, green) in the mouse kidney (left) and TCF4 immunohistochemistry in the human kidney (right, data from the Human Protein Atlas). Note the intense labeling in MD cells (arrows).

**(B)** Representative fluorescence images (left) and quantification (right) of Wnt-GFP reporter activity (top, green, from mice double transgenic for nuclear TCF/Lef:H2B-GFP reporter), and Axin2 mRNA expression (center, red) in kidney sections of WT, MD-Wnt^gof^ and MD-Wnt^lof^ mice. Note the intense labeling in MD cells (arrows). G: glomerulus. Summary of systemic and renal parameters (bottom) in the various MD-Wnt mouse models.

**(C)** Representative immunofluorescence images (left) and quantification (right) of Meis2^+^ (red, 3D projection images, top) and renin^+^ (red, bottom) cell number in kidney sections of WT, MD-Wnt^gof^ and MD-Wnt^lof^ mice. Nuclei are labeled blue with DAPI. Ns: not significant, *: p<0.05-****p<0.0001, data are mean ± SEM, n=5 glomeruli averaged for n=4 mice each group.

**Supplement Figure 3. Composition of the conditioned MD**^**geo**^ **cell culture medium and the effects of MD biologicals on kidney tissue fibrosis**.

**(A)** Mass spectrometry (left) and CCN1 ELISA (right) analysis of the composition of the conditioned MD^geo^ cell culture medium. The mass spectrum plot is representing the detected MD-derived secreted proteins in the MD^geo^ cell culture medium as indicated. LS: low-salt diet.

**(B-C)** Representative images of Picrosirius Red-stained histological sections (B) and histological analysis of interstitial fibrosis (C) in kidney tissues harvested at the end of treatment in all treatment groups as shown in Fig. 5D-F. Interstitial fibrosis was evaluated blindly by measuring the density of staining on histological sections using Image J. Scale bar is 50 μm for all panels. Ns=not significant, *P<0.05, **P<0.01, ***P<0.001, ****P<0.0001, data points represent the average of n=20 glomeruli per animal, n=6 mice each.

