## Supplementary figures and images for "Physiological activation of the nephron central command drives endogenous kidney tissue regeneration"

### Supplement Figures 1-3

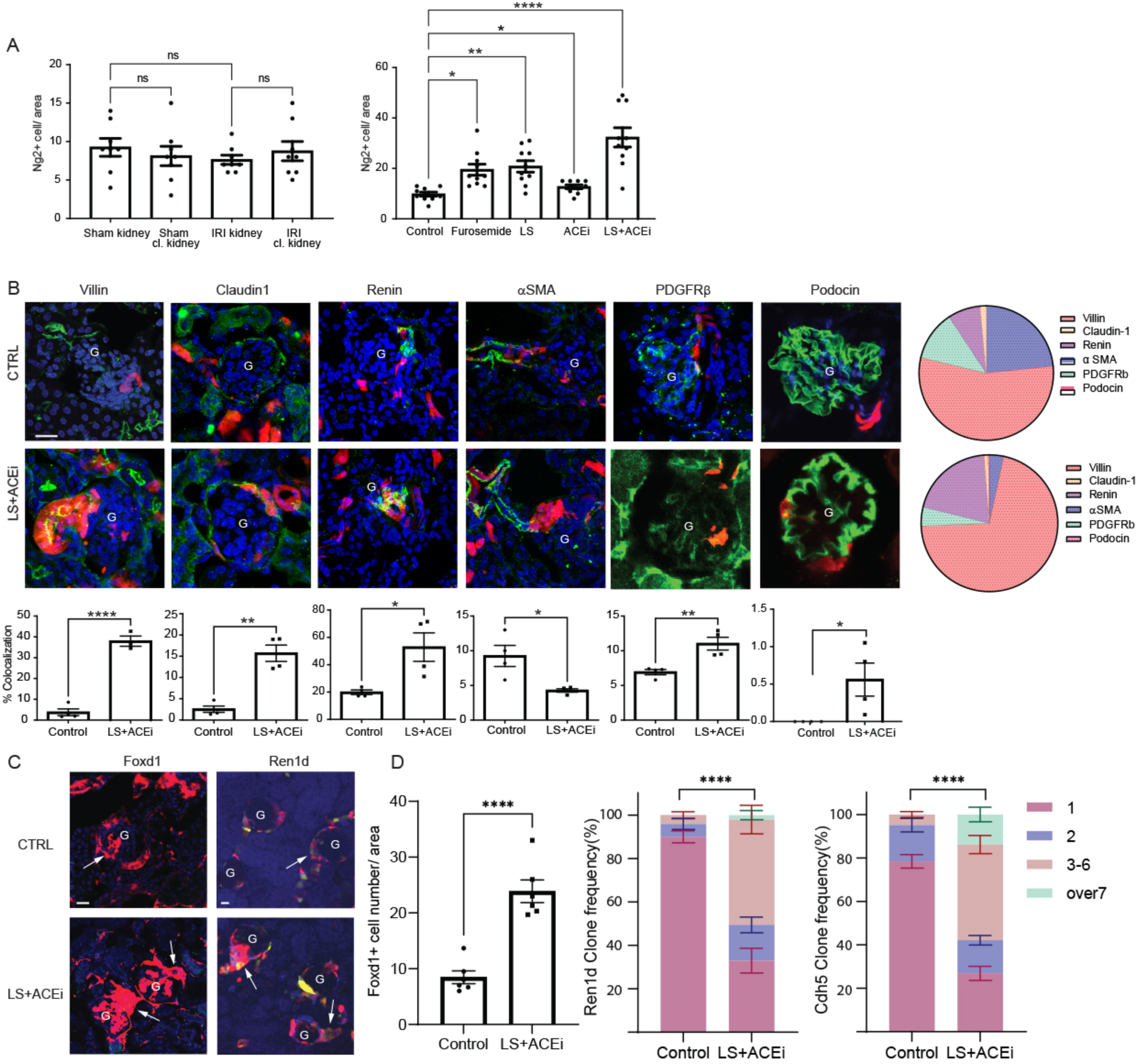

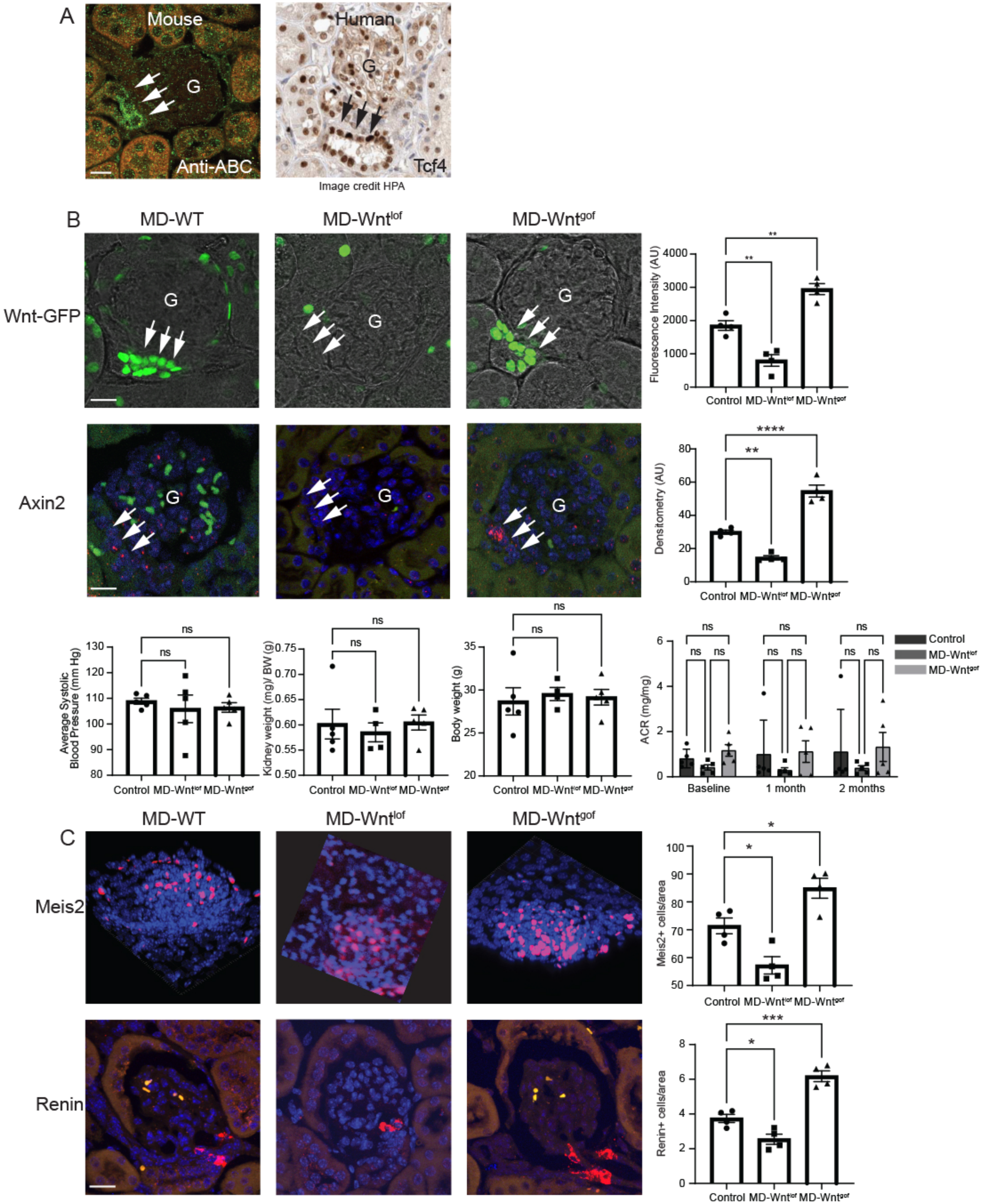

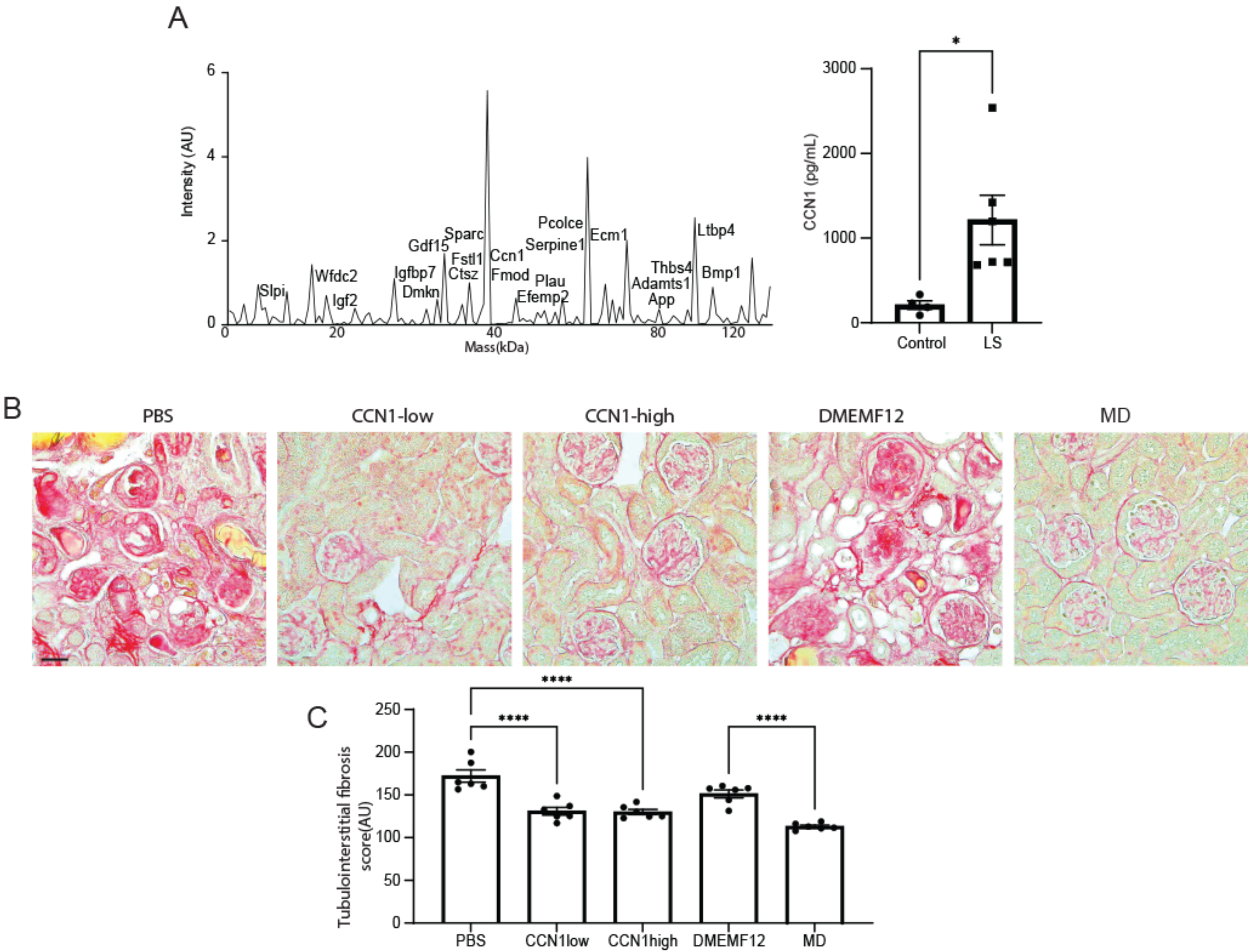
